# Quantifying predictability of gene expression from histology image

**DOI:** 10.1101/2025.11.04.686651

**Authors:** Chen-Rui Xia, Jia-Wen Yao, Ge Gao

**Author notes:** These authors contributed equally to this work (Chen-Rui Xia, Jia-Wen Yao). Authors to whom correspondence should be addressed (for G.G.).

## Abstract

Histopathological images are indispensable in clinical diagnosis, yet provide limited insight into underlying molecular states. Numerous computational models attempt to predict gene expression from histopathological images. However, a fundamental question remains unresolved: which genes can be accurately predicted and which cannot. Here, we introduce Expression Predictability Score (EPS), a metric that quantifies the predictability of each gene from images through expression-image mutual information. Empirical analyses across more than 500 slices further reveal consistent sets of highly predictable and unpredictable genes, as well as their underlying association with the physicochemical nature of H&E staining.

## Main

Histopathological imaging and spatial transcriptome technologies are complementary in terms of molecular information as well as speed and cost-effectiveness^1,2^. Thus, efforts have been made to integrate omics data with histology images^3^, and, specifically, inferring transcriptomic profiles from imaging data^4,5,6,7,8^. However, both the results of these works and independent benchmarking studies^9^ have revealed that the predictive accuracy of models varies markedly across genes, ranging from as low as 0.1 to as high as 0.9. This raises a critical question: which genes can be reliably predicted, and which results can we trust?

From an information-theoretic perspective, the mutual information between modalities (e.g., transcriptomic profiles vs histology images) defines the theoretical upper bound of cross-modal predictive performance^10,11^. Thus, we propose the Expression Predictability Score (EPS), a metric quantifying predictability of gene expression from histology image via the mutual information l(X; Y) (Methods). Mathematically, EPS is defined as the negative logarithm of graph Laplacian quadratic form calculated from the expression of gene *a* within the image-embedding kNN graph (Fig. 1a, Methods), and could be interpreted intuitively as a quantitative metric for the expression consistency of gene a across multiple regions with similar histology. We further define the Slice Predictability Score (SPS) as the average EPS across all genes within a given tissue section to quantify the slice-level predictability (Fig. 1a).

**Fig. 1:**
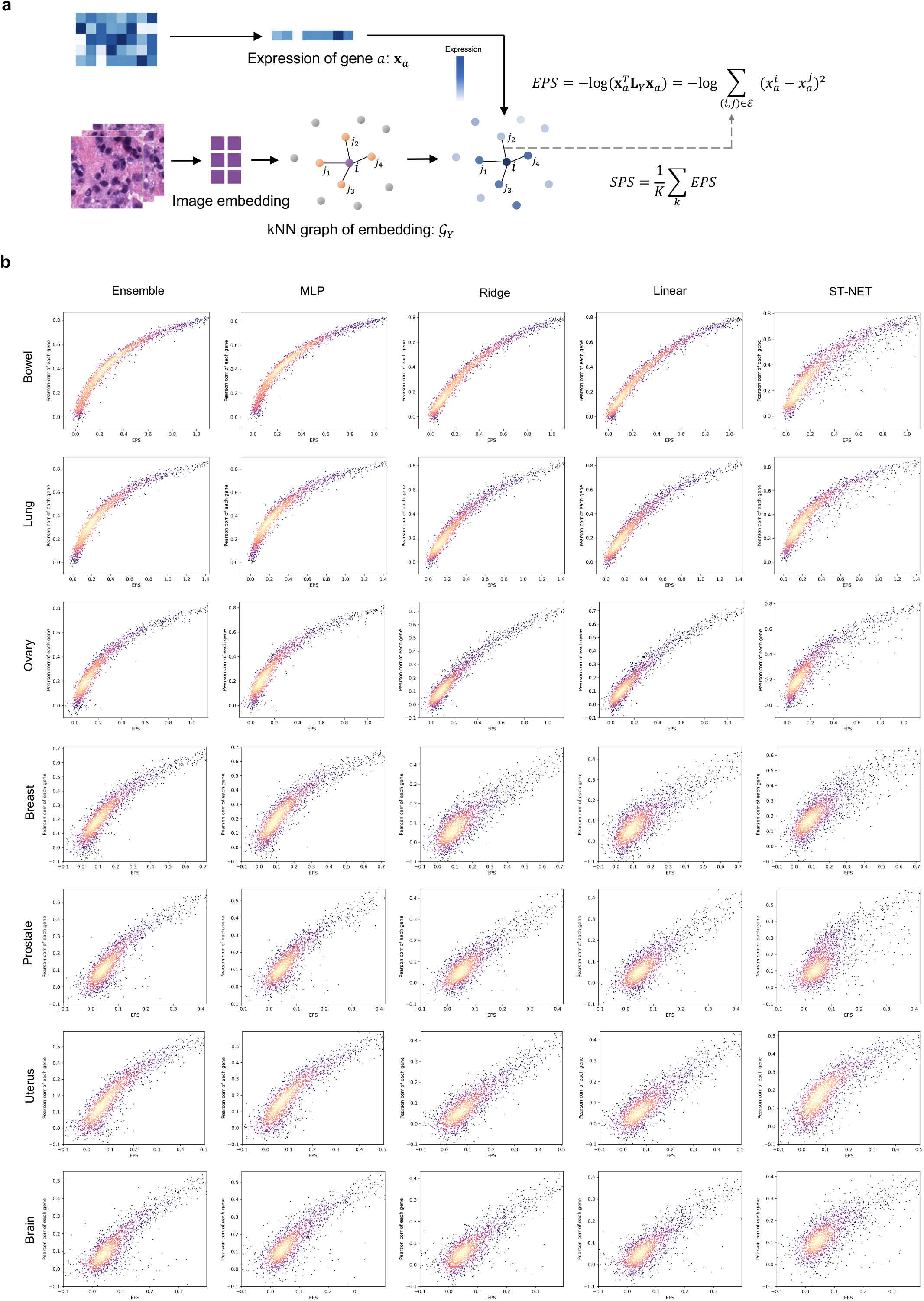
EPS quantifies gene expression predictability from histology images accurately. **a**, Schematic of the EPS and SPS calculation. For each histology image of spots/cells, we first applied an image embedding method to obtain their embeddings, and then constructed a kNN graph 𝒢_*Y*_ based on the embeddings. For a given gene *a*, its expression value (**x**_*a*_) was assigned as a node attribute on 𝒢_*Y*_. EPS of gene *a* was defined as the negative logarithm of graph Laplacian quadratic form of 𝒢_*Y*_. **L**_*Y*_ denotes the graph Laplacian matrix of 𝒢_*Y*_ (**Methods**). SPS was defined as the average EPS across all genes within a slice, *K* denotes the total gene number. **b**, Scatter plots showing the relationship between EPS (x-axis) and prediction performance (Pearson correlation, y-axis) for individual genes. Columns correspond to different prediction methods, and rows represent slices from different tissues. Points are colored according to local density.

In efforts to validate these metrics, we first assess the relationship between EPS and gene-level predictive accuracy of different models across seven tissues, including brain, bowel, lung, breast, ovary, prostate, and uterus. We carefully selected five representative models, including simple linear approaches such as linear regression and ridge regression, complex nonlinear methods such as multilayer perceptrons (denoted as MLP) and gradient-boosted trees (denoted as Ensemble), as well as the state-of-the-art deep learning–based models ST-NET^4^ (Methods). The results showed that EPS consistently quantifies the predictability, independent of tissue type and model (Fig. 1b, Extended Data Fig. 1–2 and Supplementary Table 1). Cross-model comparisons showed that all models performed poorly on genes with low EPS (i.e. low mutual information). Of note, although all models performed well in predicting high EPS genes (i.e. high mutual information), complex models—such as ST-Net, Ensemble, and MLP—achieved higher accuracy for genes with intermediate EPS (Fig. 1b, Extended Data Fig. 3). This observation suggests that overall model performance may be determined by its ability to predicting intermediate–mutual-information genes, highlighting a potential avenue for future model improvement.

Next, we systematically investigated the relationship between genes’ EPS (i.e. predictability) and properties using the HEST dataset which is consist of 510 slices covering XX (number) human tissues and. We computed the average EPS for each gene across all slices and identified top 5% vs bottom 5% predictable genes (Extended Data Fig. 4a, Supplementary Table 2). Gene Ontology (GO) enrichment analysis showed that these highly predictable genes were significantly enriched in extracellular matrix while unpredictable genes were more likely to be associated with biological membranes (Fig. 2a and 2b). We propose that such bias could be attributed to the technological nature of current H&E staining: eosin strongly binds to positively charged molecules such as collagen, which is lysine enriched, enabling strong staining of these extracellular structures^1^. In contrast, both hematoxylin and eosin are hydrophilic dyes, and therefore poorly stain hydrophobic, lipid-rich structures such as biological membranes, leading to the low predictability of membrane-associated proteins. Of interest, we noted that gene predictability exhibits tissue specificity (Fig. 2c, Extended Data Fig. 4b, also see Supplementary Table 3), which may contribute to the well-documented challenge for cross-tissue prediction^3^ and highlights the necessity of training models over datasets with comprehensive tissues covered.

**Fig. 2:**
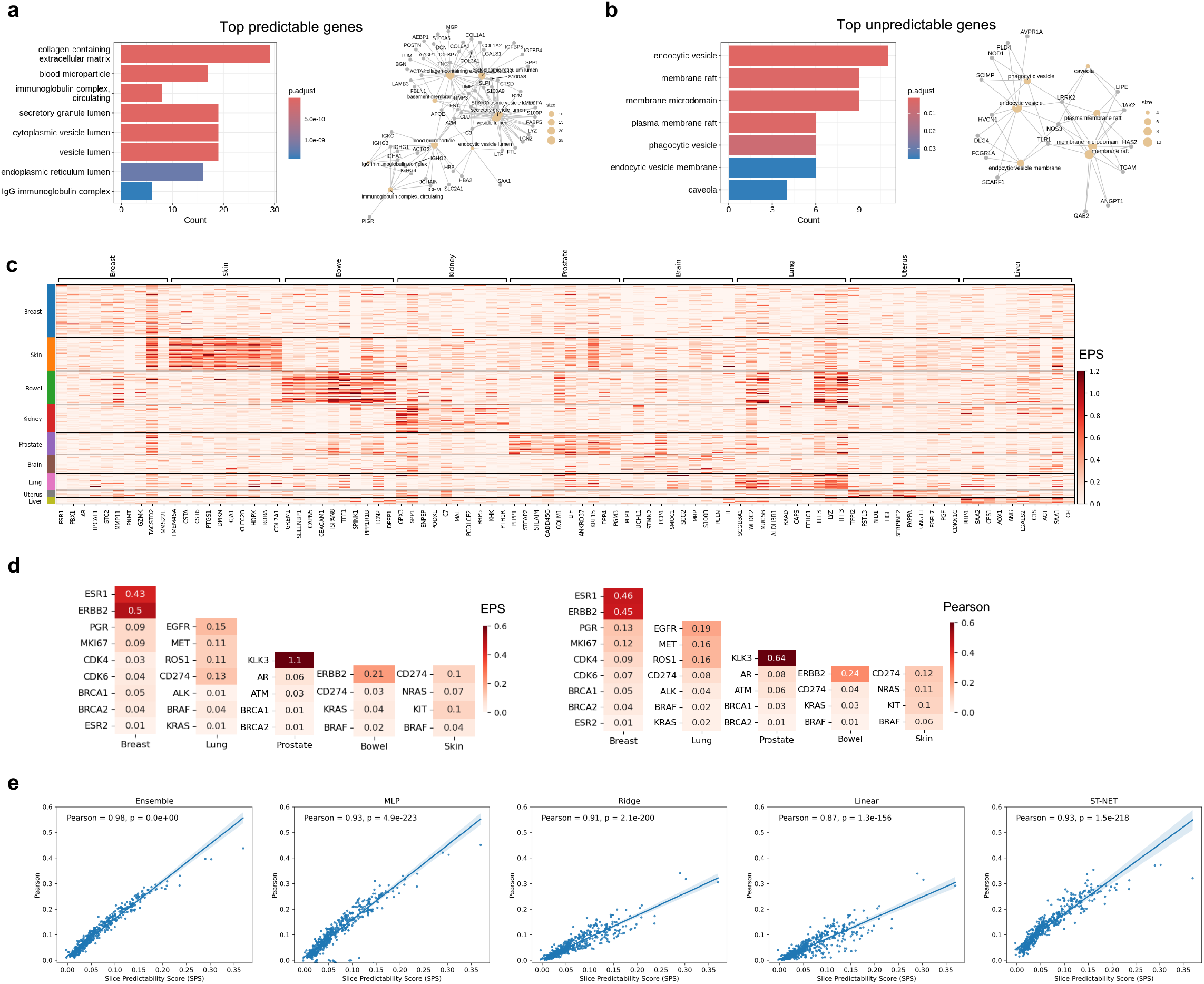
EPS and SPS analysis on HEST dataset. **a-b**, Gene Ontology Cellular Component (CC) analysis for the top predictable genes (**a**) and top unpredictable genes (**b**). Left panel: barplot of enriched GO terms in the Cellular Component category. Right panel: shows the genes and pathways in the form of a graph. **c**, Heatmap of genes with the highest EPS across different tissues. **d**, Heatmaps of EPS (left panel) and prediction performance (Pearson correlation, right panel) for known biomarkers across multiple cancer types. **e**, Scatter plots showing the relationship between SPS (x-axis) and prediction performance (Pearson correlation, y-axis) across slices in the HEST dataset.

We further focused on the predictability of clinically meaningful genes in cancer^12^. Encouragingly, we found that *PSA* (*KLK3*) in prostate cancer, as well as *HER2* (*ERBB2*) and *ESR1* in breast cancer, exhibited high predictability (high EPS), and correspondingly strong predictive performance (Fig. 2d). However, some other clinically important markers, such as *PD-L1* (*CD274*), *BRCA1*, and *BRCA2* showed low predictability and poor predictive performance (Fig. 2d), suggesting that the histopathological imaging-based inference should be interpreted with caution.

Consistently, we also found the slice-level metric SPS is strongly correlated with slice-level prediction accuracy of all models (Fig. 2e and Extended Data Fig. 4c). While model performance varied on high-SPS slices, all models showed similarly poor performance on low-SPS slices (Fig. 2a, Extended Data Fig. 4d). These findings reflect the lack of mutual information that prevents any model from making accurate predictions, further confirming the decisive role of mutual information in expression predictability.

Of note, EPS and SPS are not limited to image–expression data and can be applied to any paired multi-omics datasets. For example, using multi-omics data from the NeurIPS 2021 Open Problem Competition^13^, we employed EPS and SPS to evaluate the predictability of ATAC to RNA, as well as surface protein to RNA. They also demonstrated consistently strong performance (Extended Data Fig.5-6), highlighting their potential as general indicators of cross-modal predictability.

Owing to the intrinsic complexity of the image distribution, we currently employ the published image model^14^ when inferring the kNN graph for EPS calculation (**Methods**).Thus, a suboptimal embedding may introduce noise into the graph structure and further lead to an underestimation of EPS. However, we believe that the rapid development of image models would effectively alleviate this problem.

In summary, we have developed the Expression Predictability Score (EPS), a metric that quantifies the predictability of genes from histopathological images. By analyzing a large number of slices, we found that the bias in gene predictability may be attributed to the hydrophilic nature of H&E dyes, and EPS can also serve as a quantitative metric for guiding the development of new tissue-staining methods to enable accurate prediction of specific genes or biomarkers. We recommend that EPS be carefully considered for each gene prior to interpreting the results of any image-based gene expression prediction model. The full code is publicly available at https://github.com/gao-lab/EPS.

## Methods

### Expression Predictability Score

For spatial omics data with *N* spots, let **x**_*a*_ denote the scaled expression of gene *a*, and *Y* represents the distribution of the paired histology images. Given to the complexity of the image distribution *Y*, we here introduce a spot-wise k-nearest neighbor (kNN) graph *𝒢*_*Y*_: derived from pretained image foundation model (here is the state-of-the-art UNI model^14^, with *k*=5 by default). Then,for gene *a*, EPS could be represented as the negative logarithm of the graph Laplacian quadratic form of **x**_*a*_ on *𝒢*_*Y*_:

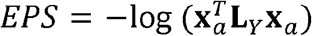

where **L**_*Y*_ denotes the graph Laplacian matrix corresponding to the graph *𝒢*_*Y*_:

Now we’d derive the relationship between EPS and the mutual information *I*(**x**_*a*_; *Y*) analytically. Specifically, we express the mutual information in terms of conditional entropy:

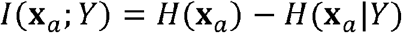

Given the fact the *H*(**x**_*a*_) is a constant, we only need to consider *H*(**x**_*a*_|*Y*). However, the analytical estimation of conditional distribution *H*(**x**_*a*_|*Y*) is not trival. Therefore, we use the Gaussian Markov Random Field (GMRF) probabilistic model^15^ to link the spots’ gene expression (Gaussian distribution) with their histological images (in the format of graph). In practice, we assume that **x**_*a*_ follows a conditional distribution over *𝒢*_*Y*_:

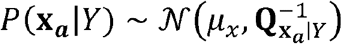

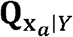 is the precision matrix (the inverse of the covariance matrix) of the GMRF, and 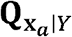 is proportional to the graph Laplacian matrix **L**_*Y*_ of *𝒢*_*Y*_^15^;

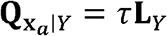

Tthe maximum likelihood of τ could be given by (see Supplementary Note 1 for details):

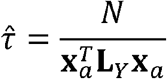

*N* is the spot number in the slice.

The entropy of GMRF *P*(**x**_*a*_|*Y*) is^15^:

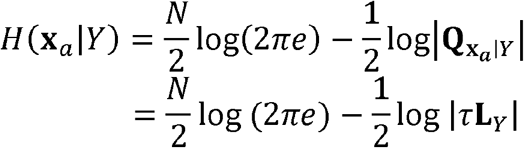

For the N-order matrix **L**_*Y*_, we have 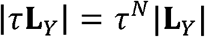 thus:

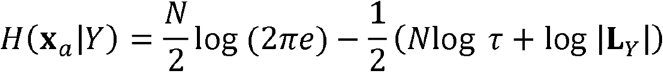

So, the mutual information could be rewritten as:

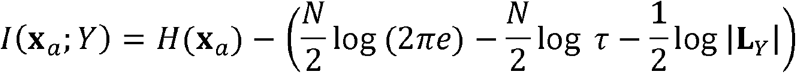

We omit all of constant values, including 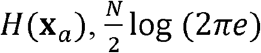, and 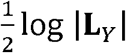, and obtain:

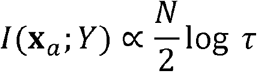

Substituting the maximum likelihood estimate 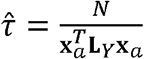, and omitting constant terms, we obtain:

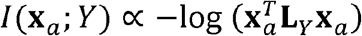

i.e.:

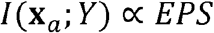

### Data collection and pre-process

Seven 10x spatial slices were obtained from the official 10x Genomics website. These datasets encompass tissues from diverse organs and disease states, each paired with a corresponding H&E-stained image (Supplementary Table 1). The HEST data were obtained from the HEST dataset^16^, from which we selected human slices and excluded slices containing fewer than 100 spots, leaving a total of 510 slices. All spatial slices were processed using the same pipeline. For H&E images, we first scaled the images to 0.5□μm per pixel (mpp), and then extracted square tiles (224 × 224 pixels, 112□μm in diameter) centered at each spot coordinate. Then, image of each tile was embedded into a 1024-dimensional vector using the pretrained UNI^14^ model. For gene expression data, we selected the top 2,000 highly variable genes shared across all slices, to ensure fair cross-slice comparisons. The expression data were then normalized, log-transformed (log1p), and scaled following the standard Scanpy workflow^17^.

Single-cell multi-omics datasets were obtained from the NeurIPS 2021 Open Problem competition^13^ and include two data types: 10x Multiome (RNA and ATAC) and CITE-seq (RNA and surface protein). The 10x Multiome dataset comprises 13 distinct samples (batches), while CITE-seq comprises 12 batches. For RNA data, we applied the same workflow as described above, including selection of the top 2,000 highly variable genes, normalization, log1p transformation, and scaling. For surface protein data, raw counts were log-normalized, scaled, and then use principal component analysis (PCA) to reduce dimensionality to 20 components. For ATAC data, dimensionality reduction was performed using the spectral decomposition method described in SnapATAC2^18^ with default parameters, yielding 30 components.

### Prediction models

We constructed a variety of baseline models to predict gene expression from histological images:

1. Linear regression (denoted as Linear), implemented using scikit-learn.
2. Ridge regression (denoted as Ridge), also implemented in scikit-learn, with *α*= 1.
3. Multilayer perceptron regression (denoted as MLP), implemented in PyTorch. Specifically, the MLP consisted of a hidden layer with 256 units and ReLU activation. The mean squared error (MSE) was used as the loss function. Models were trained for 100 epochs using the AdamW optimizer, with a learning rate of 1 ×10□ ^3^ and a batch size of 128.
4. Gradient-boosted trees (denoted as Ensemble), implemented using LightGBM with 100 trees, trained separately for each gene.

For each tissue slice, data were randomly split into training and test sets at an 80:20 ratio. In addition, we included the previously published ST-Net^4^ model, trained using the same preprocessing and train/test splits as the baseline models and following the training procedures described by the original authors. For single-cell multi-omics data, we trained only the MLP as the baseline model, as it was the most commonly choice among the winning teams in the original competition^13^.

Model performance on the test set was evaluated using Pearson correlation, Spearman correlation, and Root Mean Square Error (RMSE). Then, the sample-level metric was calculated as the average across all genes.

### Statistical analysis

Pearson and Spearman correlation coefficients, as well as the corresponding *P*-values, were calculated using the “scipy.stats” Python package (v1.16.1). Gene Ontology (GO) enrichment analysis was performed using the “clusterProfiler” R package^19^ (v4.6.2), with a *P-*value threshold of 0.05. *P*-values were adjusted using the Benjamini–Hochberg method.

## Data availability statement

All datasets used in this study were already published and were obtained from public data repositories. 10x spatial slices used in this study are recoded in Supplementary Table 1, including downloading URLs. HEST dataset^16^ is available at Hugging Face (https://huggingface.co/datasets/MahmoodLab/hest). NeurIPS 10x Multiome and CITE-seq data^13^ is available at GSE194122 (https://www.ncbi.nlm.nih.gov/geo/query/acc.cgi?acc=GSE194122).

## Code availability statement

The source code of calculating EPS and SPS, as well as codes to reproduce the results in this paper, can be accessed from https://github.com/gao-lab/EPS under MIT license.

## Acknowledgments

We thank for Drs. Z. Cao, Z. Zhang, F. Tang, and X.S. Xie at Peking University for their helpful discussions and comments during the study. This work was supported by funds from the National Key Research and Development Program of China (2022ZD0115004), as well as the State Key Laboratory of Gene Function and Modulation Research, the Beijing Advanced Innovation Center for Genomics (ICG) at Peking University, the Changping Laboratory, and the Shaw Foundation Hong Kong Limited. The research of C.R.X. is supported in part by the National Natural Science Foundation of China (grant no. 323B2017).

## Author contributions

G.G. conceived the study and supervised the research. C.R.X. designed and implemented the computational framework. C.R.X. and J.W.Y conducted experiments with guidance from G.G. C.R.X., J.W.Y., and G.G. wrote the manuscript.

## Competing interests

The authors declare that they have no competing interests.

## Figure legends

**Extended Data Fig. 1:**
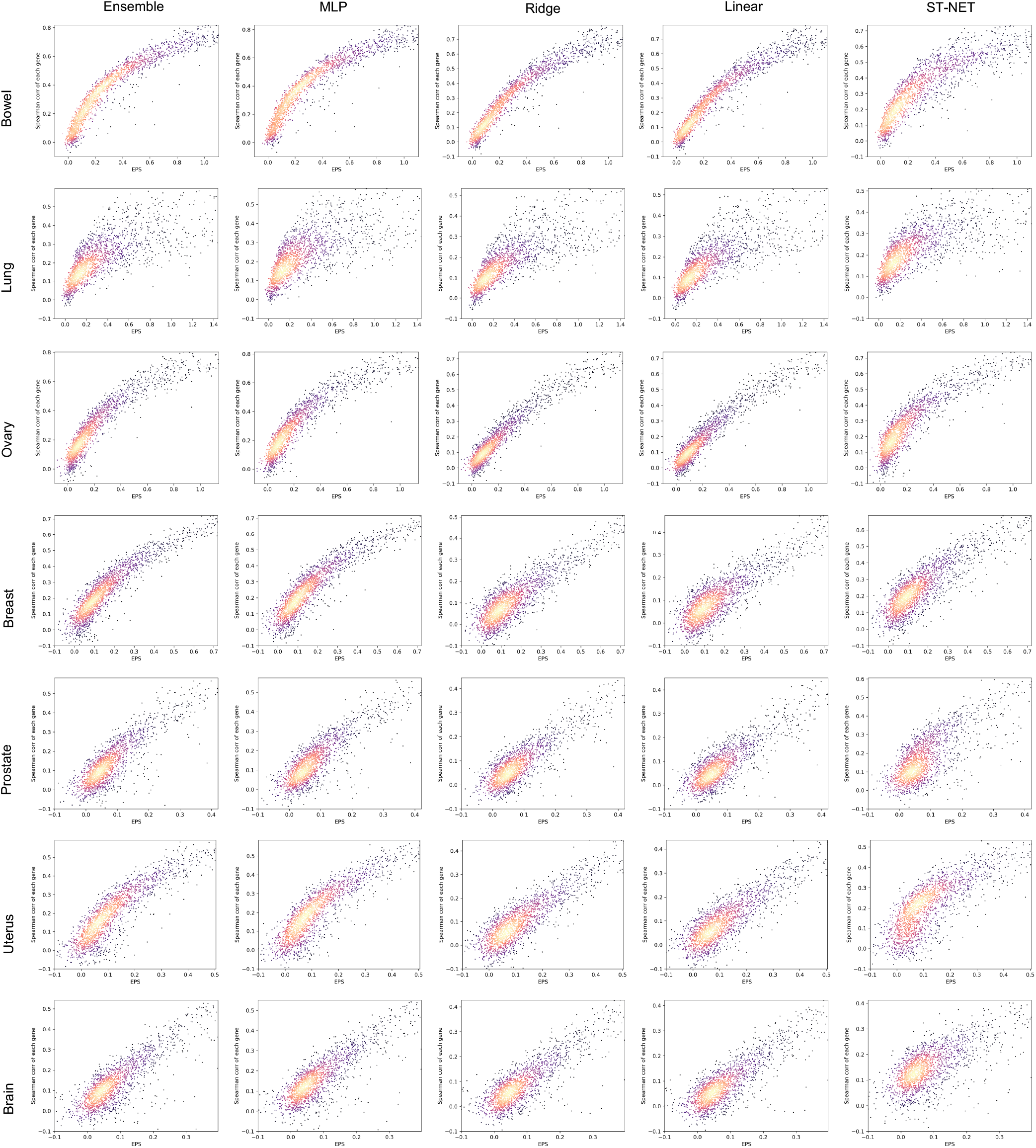
Correlation between EPS and Spearman correlation. Scatter plots showing the relationship between EPS (x-axis) and prediction performance (Spearman correlation, y-axis) for individual genes. Columns represent different prediction methods, and rows correspond to slices from different tissues. Points are colored according to local density.

**Extended Data Fig. 2:**
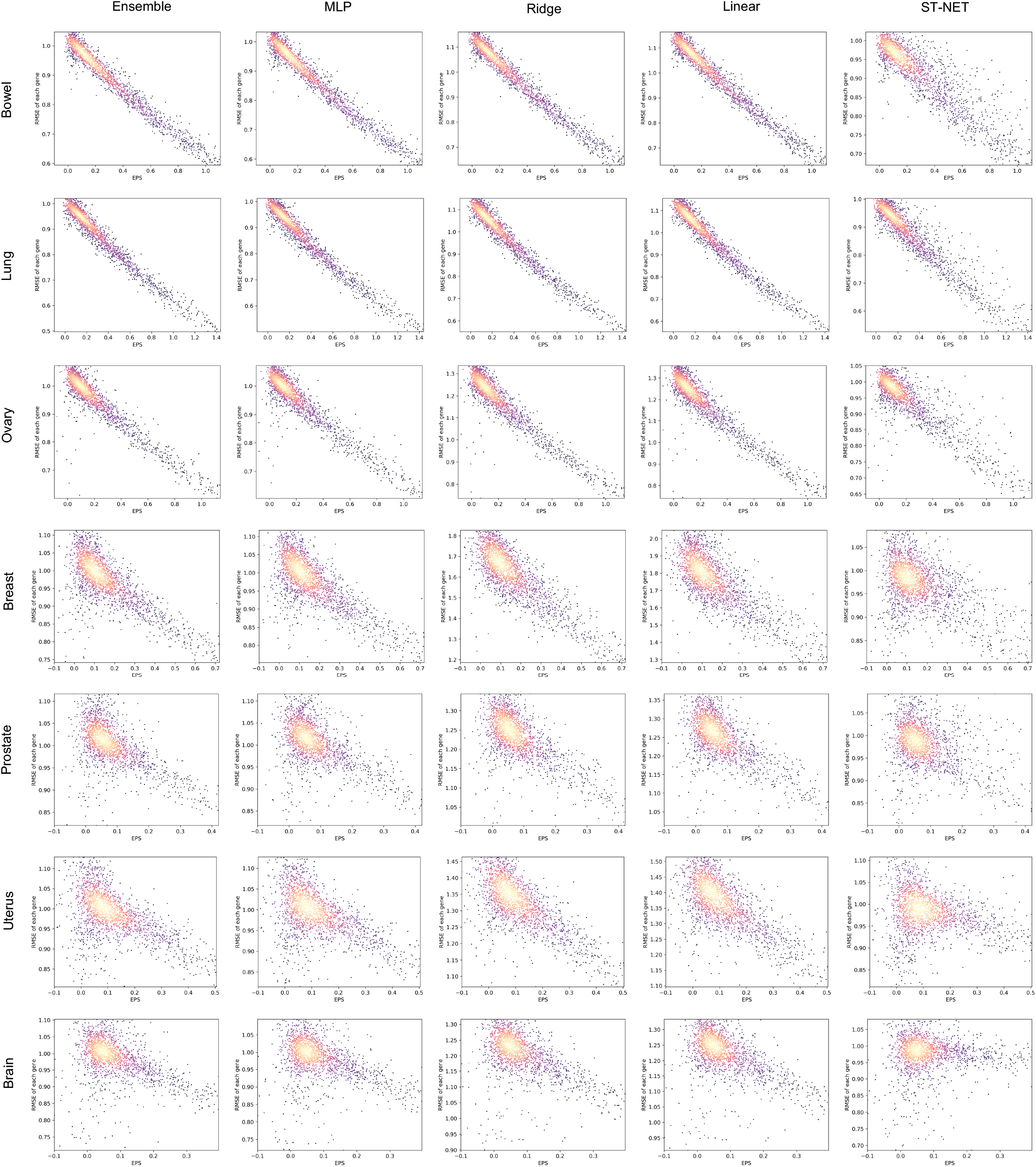
Correlation between EPS and RMSE. Scatter plots showing the relationship between EPS (x-axis) and prediction error (RMSE, y-axis) for individual genes. Columns represent different prediction methods, and rows correspond to slices from different tissues. Points are colored according to local density.

**Extended Data Fig. 3:**
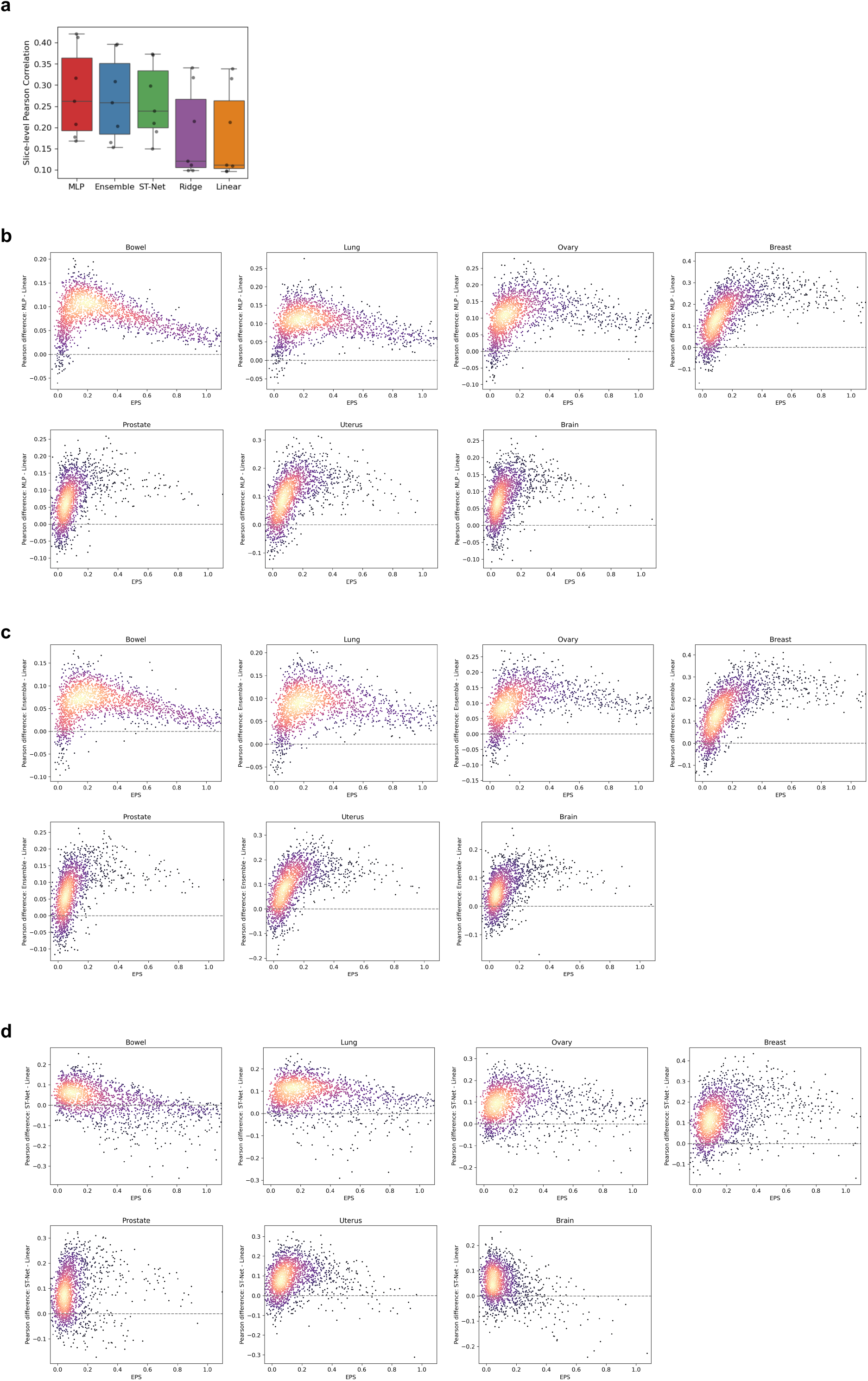
Comparing prediction accuracy of prediction methods in 10x slices. **a**, Boxplot showing the average per-slice prediction performance (Pearson correlation) of each method across seven 10x slices. **b-d**, Scatter plots showing the per-gene performance differences between MLP (**b**), Ensemble (**c**), and ST-NET (**d**) relative to the Linear model in 10x slices. Points are colored according to local density.

**Extended Data Fig. 4:**
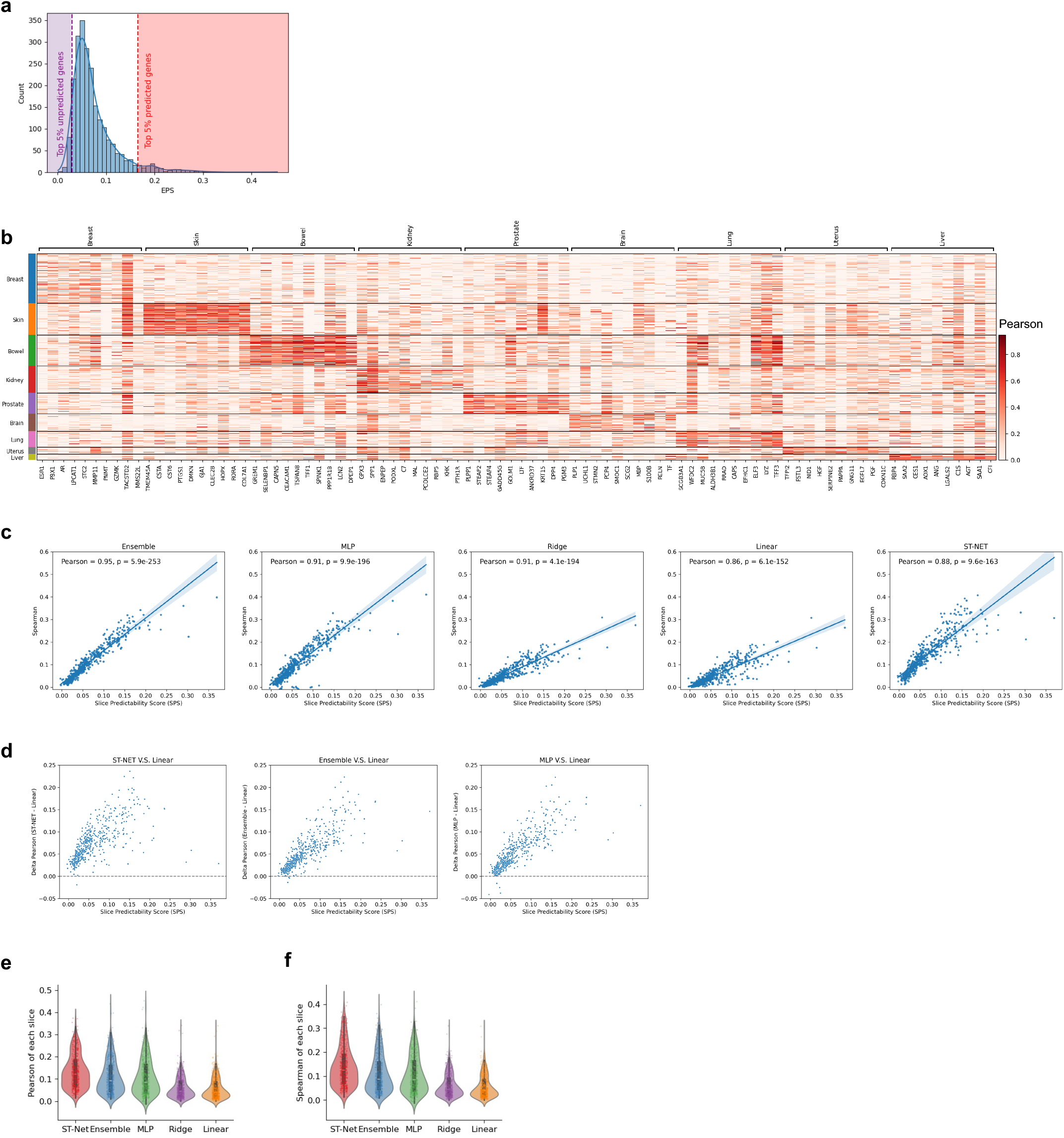
Comparison of prediction accuracy across methods in the HEST dataset. **a**, Histogram showing the distribution of average EPS across slices in the HEST dataset. Purple and red shadows indicate the “top predictable genes” and “top unpredictable genes”, respectively. **b**, Heatmap of prediction performance (Pearson correlation) for genes with the highest EPS across different tissues (as shown in Fig. 2d). **c**, Scatter plots showing the relationship between SPS (x-axis) and prediction performance (Pearson correlation, y-axis) across slices in the HEST dataset. **d**, Scatter plots showing the per-slice performance differences of ST-NET, Ensemble and MLP relative to the Linear model in the HEST dataset. **e-f**, Violin plots showing the distributions of Pearson correlation (**e**) and Spearman correlation (**f**) for each method in the HEST dataset.

**Extended Data Fig. 5:**
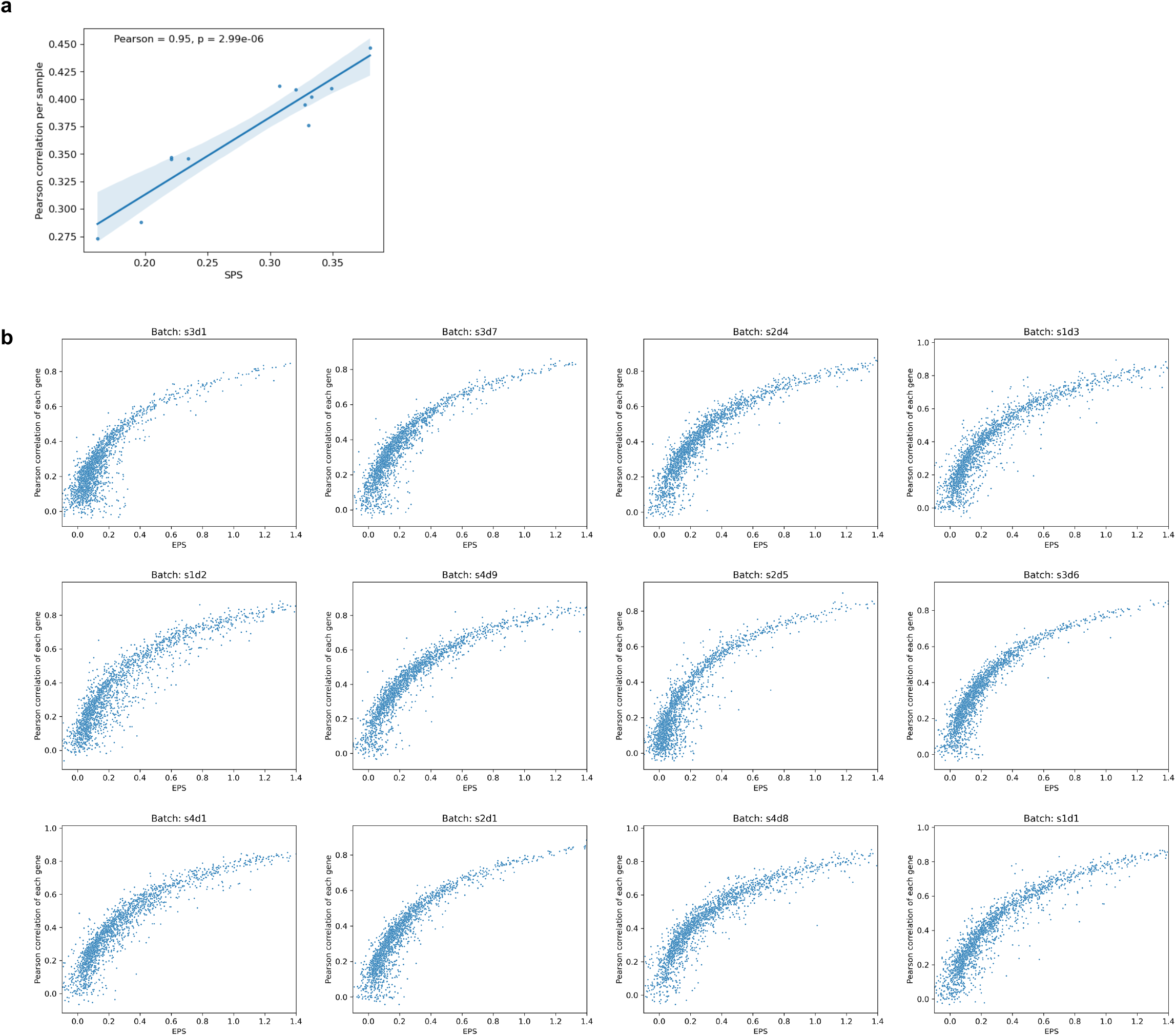
EPS and SPS results in CITE-seq data. **a**, Scatter plot showing the relationship between SPS (x-axis) and prediction performance (Pearson correlation, y-axis) in each batch. **b**, Scatter plots showing the relationship between EPS (x-axis) and prediction performance (Pearson correlation, y-axis) for individual genes, each panel represents a batch.

**Extended Data Fig. 6:**
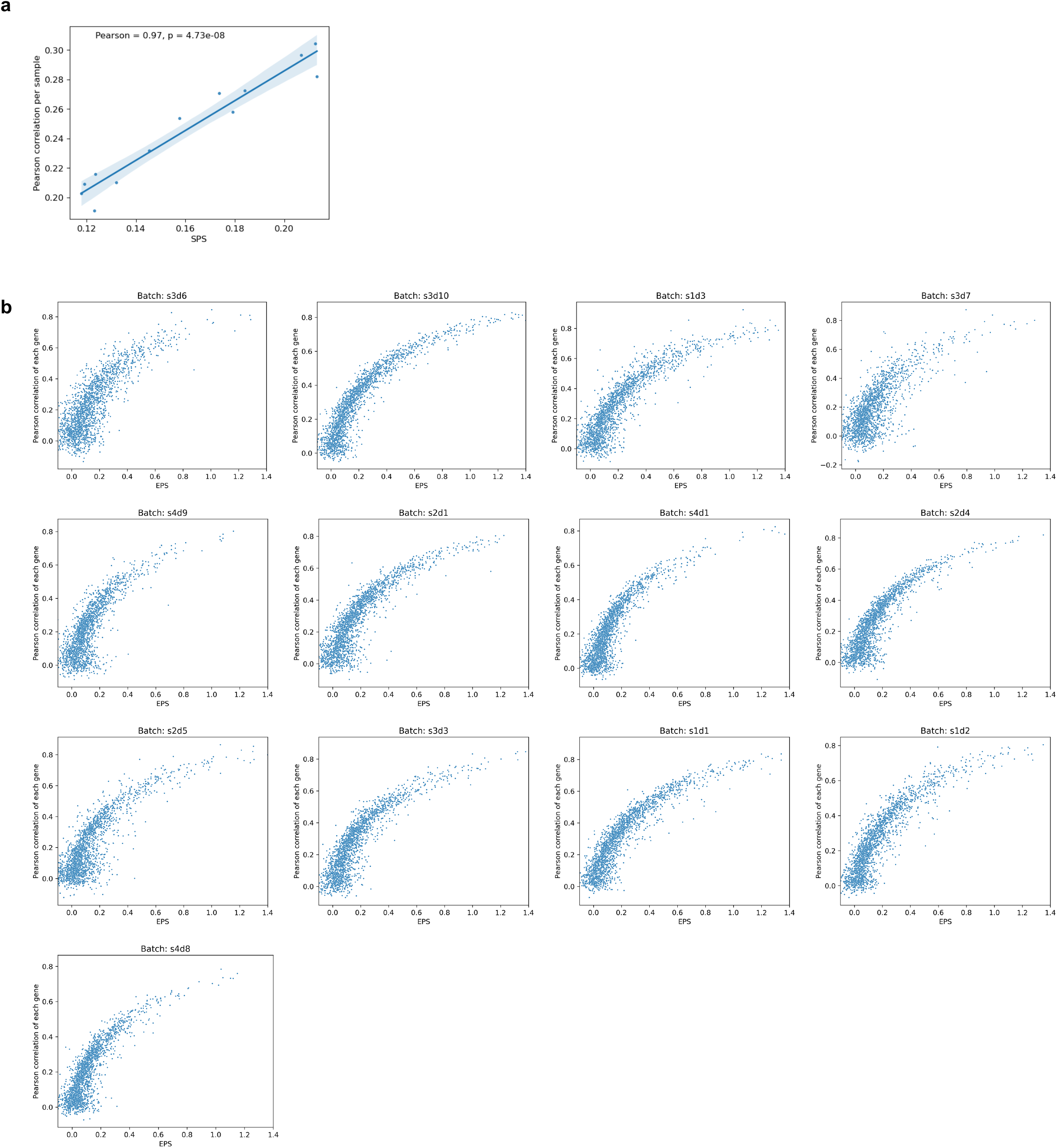
EPS and SPS results in 10x Multiome data. **a**, Scatter plot showing the relationship between SPS (x-axis) and prediction performance (Pearson correlation, y-axis) in each batch. **b**, Scatter plots showing the relationship between EPS (x-axis) and prediction performance (Pearson correlation, y-axis) for individual genes, each panel represents a batch.

## Notes

https://github.com/gao-lab/EPS

### Competing Interest Statement

The authors have declared no competing interest.

