## Supplementary Note 1 for "Quantifying predictability of gene expression from histology image"

**SUPPLEMENTARY** **MATERIALS**

**Supplementary Note 1**

For the Gaussian Markov Random Field (GMRF)：

$$P\left( \mathbf{x}_{\boldsymbol{a}} \right|Y)\sim\mathcal{N}\left( \mu_{x},\mathbf{Q}_{\mathbf{x}_{\boldsymbol{a}}\boldsymbol{\mid}Y}^{\boldsymbol{-1}} \right)$$

$\mathbf{Q}_{\mathbf{x}_{\boldsymbol{a}}\boldsymbol{\mid}Y}$ is the precision matrix, and $\mathbf{Q}_{\mathbf{x}_{\boldsymbol{a}}\boldsymbol{\mid}Y}=\tau\mathbf{L}_{Y}$. Here, we use maximum likelihood estimation (MLE) to find the optimal estimate of $\hat{\tau}$. Since $\mathbf{x}_{a}$ is 1-D Gaussian distribution (with the mean of 0 and a variance of 1), the log-likelihood function is a strictly convex function, so the MLE can find a strict global maximum rather than saddle point. The likelihood function of $P\left( \mathbf{x}_{\boldsymbol{a}} \right|Y)$ is:

$$L\left( \tau;\mathbf{x}_{a},\mathbf{L}_{Y} \right)=\frac{\left| \mathbf{Q}_{\mathbf{x}_{\boldsymbol{a}}\boldsymbol{\mid}Y} \right|^{1/2}}{2\pi} \exp\left( -\frac{1}{2}\mathbf{x}_{a}^{T}\mathbf{Q}_{\mathbf{x}_{\boldsymbol{a}}\boldsymbol{\mid}Y}\mathbf{x}_{a} \right)$$

$$=\frac{\left| \tau\mathbf{L}_{Y} \right|^{1/2}}{2\pi} \exp\left( -\frac{\tau}{2}\mathbf{x}_{a}^{T}\mathbf{L}_{Y}\mathbf{x}_{a} \right)$$

For computational convenience, we maximize the log-likelihood function $\mathcal{l(}\tau)=\log L(\tau)$：

$$\mathcal{l}\left( \tau\right)=-\log2\pi+\frac{1}{2}\log\left| \tau\mathbf{L}_{Y} \right|-\frac{\tau}{2}\mathbf{x}_{a}^{T}\mathbf{L}_{Y}\mathbf{x}_{a}$$

For the N-order matrix $L_{Y},$ we have $\left| \tau L_{Y} \right|=\tau^{N}|L_{Y}|$, thus:

$$\log\left| \tau\mathbf{L}_{Y} \right|=\log\left( \tau^{N} \right)+\log\left| \mathbf{L}_{Y} \right|=Nlog\tau+log\left| \mathbf{L}_{Y} \right|$$

Therefore, the log-likelihood function can be simplified as:

$$\mathcal{l(}\tau)=-\log2\pi+\frac{N}{2}\log\tau+\frac{1}{2}log\left| \mathbf{L}_{Y} \right|-\frac{\tau}{2}\mathbf{x}_{a}^{T}\mathbf{L}_{Y}\mathbf{x}_{a}$$

We next take the derivative of $\tau$ and set it to zero in order to obtain the maximum likelihood estimate $\hat{\tau}$ that maximizes $\mathcal{l(}\tau)$:

$$\frac{\partial\mathcal{l}}{\partial\tau}=\frac{N}{2\tau}-\frac{1}{2}\mathbf{x}_{a}^{T}\mathbf{L}_{Y}\mathbf{x}_{a}=0$$

Solving this equation, we get the maximum likelihood estimate of $\hat{\tau}$：

$$\hat{\tau}=\frac{N}{\mathbf{x}_{a}^{T}\mathbf{L}_{Y}\mathbf{x}_{a}}$$
